# Distribution map of peristaltic waves in the chicken embryonic gut reveals importance of ENS and inter-region cross talks along the gut axis

**DOI:** 10.1101/2021.12.08.471708

**Authors:** Yuuki Shikaya, Yuta Takase, Ryosuke Tadokoro, Ryo Nakamura, Masafumi Inaba, Yoshiko Takahashi

## Abstract

Gut peristaltic movements recognized as the wave-like propagation of a local contraction are crucial for effective transportation and digestion/absorption of ingested materials. Although the physiology of gut peristalsis has been well studied in adults, it remains largely unexplored how the cellular functions underlying these coordinated tissue movements are established along the rostral-caudal gut axis during development. The chicken embryonic gut serves as an excellent experimental model for elucidating the endogenous potential and regulation of these cells since peristalsis occurs even though no ingested material is present in the moving gut. By combining video-recordings and kymography, we provide a spatial map of peristaltic movements along the entire gut posterior to the duodenum: midgut (jejunum and ileum), hindgut, caecum, and cloaca. Since the majority of waves propagate bidirectionally at least until embryonic day 12 (E12), the sites of origin of peristaltic waves (OPWs) can unambiguously be detected in the kymograph. The spatial distribution map of OPWs has revealed that OPWs become progressively confined to specific regions/zones along the gut axis during development by E12, and that such specific zones are largely conserved between different individuals implying genetic regulation for OPW determination. We have also found that the enteric nervous system (ENS) is essential for the OPW patterning since an ablation of ENS or blocking neural activity by tetrodotoxin disrupts the confined pattern of OPWs, resulting in a failure of transportation of inter-luminally injected ink. Finally, we have discovered a functional coupling of the endpoint of hindgut with the cloaca. When surgically separated, the cloaca ceases its acute contractions that would normally occur concomitantly with the peristaltic rhythm of the hindgut. Our findings shed light on the intrinsic regulations of gut peristalsis, including unprecedented ENS contribution and inter-region cross talk along the gut axis.

**Contribution to the field statement:** It has been well accepted that the gut peristalsis is important in adults, where a luminal content (ingested material) mechanically influences peristalsis. However, the endogenously regulated cellular mechanisms that initiate and sustain peristalsis have poorly been explored, and this might be a reason why few therapeutic treatments have been available for gut diseases associated with peristaltic dysfunction. Recent studies have shown that peristaltic movements occur in the embryonic gut, suggestive of genetic involvement. However, how the peristaltic movement is coordinately patterned along the gut axis remains unknown, because most studies have used isolated fragments of the gut. In our study, we examined the entire gut posterior to the duodenum of chicken embryos to produce a spatial map of peristaltic movements during gut development. Using the map of origins of peristaltic waves (OPWs), we found previously unappreciated roles of enteric nervous system for the OPW patterning and for the physiological functions of embryonic gut tissues. Furthermore, we propose that peristaltic movements might mediate inter-region crosstalk along the gut axis. Our findings provide novel insights into the mechanisms by which the gut peristalsis is established during development at the cellular basis.

## Introduction

During feeding, food is transported aborally from the mouth through the esophagus to the stomach, small and large intestines, where the ingested material is digested and absorbed. Peristaltic movements in the intestines, recognized as the wavelike propagation of a local contraction, play a crucial role in bolus transportation. Although conventional physiology has stated that peristalsis is triggered by a bolus in adults, how this is regulated at the cellular basis is not fully understood (Huizinga et al., 1998; Huizinga and Lammers, 2009; Spencer et al., 2016). Many gut-related disorders, including irritable bowel syndrome and post-operative paralytic ileus, are accompanied by peristaltic dysfunctions, resulting in severe digestive problems (Bauer and Boeckxstaens, 2004; Bornstein et al., 2004; Bouchoucha et al., 2006; Mattei and Rombeau, 2006; Pimentel et al., 2002). Thus, it is important to elucidate the cellular mechanisms by which gut peristalsis is sustained and regulated.

The embryonic chicken gut serves as an excellent model system when working toward this goal, as the gut undergoes peristalsis even before hatching (no feeding), and also because gut-constituting cells and tissues are easily manipulable experimentally. Chevalier and co-workers reported pioneering studies using chicken embryonic guts, in which they described changes in speed and frequencies of local peristaltic waves during development (Chevalier et al., 2020; Chevalier et al., 2019; Chevalier et al., 2017). They observed irregular and discontinuous contractions in early embryos at E5, which was followed by emergence of peristaltic propagations around E7. It was also shown that whereas circular muscle contractions at E12 did not require the activity of enteric nervous system (ENS) (Boeckxstaens et al.), the ENS governed the gut peristalsis by E16 (Chevalier et al., 2019; Chevalier et al., 2017). The ENS-independent contractions were also observed in mouse and zebrafish embryos (Holmberg et al., 2007; Roberts et al., 2010). Since these studies focused on the timing of local contractions-peristalsis, these data could be acquired using fragments of specific gut regions at specific times.

However, the gut is a long organ, where peristalsis occurs widely in a complex manner along the gut axis. Therefore, the spatial information of the peristaltic patterns along the entire gut axis is necessary for understanding the regulation of peristalsis. However, such information has been lacking. In this study, we have produced a “map of peristalsis” along the developing gut posterior to the duodenum, which includes jejunum, ileum, hindgut, caecum, and cloaca. Combining video recordings of non-interrupted gut specimens and kymographic analyses, we have asked four specific questions: 1) Where do the peristaltic waves originate along the entire gut axis? (i.e. where are the <u>o</u>rigins of <u>p</u>eristaltic <u>w</u>aves (OPWs) located? 2) Is the distribution of OPWs stochastic, or is it ordered by some rules? 3) Does the ENS play a role in the patterning of peristaltic waves? 4) Are the peristaltic patterns coordinated between different regions along the gut axis; and if so, how is such coordination regulated? By addressing these questions, we have found that during the development of the midgut (jejunum and ileum), randomly distributed OPWs become progressively confined to distinct zones along the gut axis, which are largely conserved among different individuals, implying that intrinsic regulations are involved. In addition, the ENS plays a role in the determination of the zones of OPWs, and also in the effective transportation of inter-luminally injected ink particles. Furthermore, the peristaltic rhythm in the hindgut is tightly coupled with acute contractions of the cloaca (anus-like structure in avians), revealing a coordination between these tissues.

## Materials and Methods

### Chicken embryos

Fertilized chicken eggs were obtained from the Yamagishi poultry farms (Wakayama, Japan). Embryos were staged according to Hamburger and Hamilton (HH) series (Hamburger and Hamilton, 1951) or embryonic day (E). All animal experiments were conducted with the ethical approval of Kyoto University (#202110).

### Motility monitoring of the gut and kymograph preparation

The gut from duodenum to cloaca was dissected from embryos, and placed into silicone-coated petri dished filled with high-glucose DMEM (Wako, 048-33575) warmed at 38.5 °C with a heating plate (MSA Factory, PH200-100/PCC100G). To avoid drifts of the specimen during imaging, gut-attached remnants such as pancreas, vitelline membrane and cloaca were pinned to the silicone dish with fine needles. Following 10 min of resting, time-lapse images were captured using the Leica MZ10 F microscope with a DS-Ri1 camera (Nikon). The obtained images were processed by ImageJ software (NIH) to analyze the gut peristalsis. The original images were converted into 8-bits images in pseudo-color to highlight the peristaltic contractions. These processed images were subjected to the Reslice of ImageJ to generate kymographs along the gut axis. The positions of the OPWs and the velocity of propagation waves were reliably deduced from the apices and slopes of the contraction lines in the kymograph, respectively. For the experiments demonstrated in Fig. 5, the medium contained 10 μM tetrodotoxin (nacalai tesque, 32775-51).

### Surgical manipulations

For the ablation of enteric nervous system, the neural tube of the vagal region (somite levels 1-7) of E1.5 (HH10) embryo was surgically removed using a sharpened tungsten needle. Separations of caeca from the main tract and of the cloaca from the hindgut were conducted using micro-scissors.

### Ink injection into the gut

A solution of highlighter ink (Takase et al., 2013) was injected into duodenum of a dissected gut. The specimens were placed on a glass bottom dish (Greiner, 627860) filled with high-glucose DMEM. Time-lapse images were captured using a Nikon A1R confocal microscope at 38.5°C. For Ca^2+^-free medium treatment, guts were soaked in Ca^2+^-free DMEM (Thermo Fisher Scientific, 21068-028), and incubated for 10 min at 38.5 °C prior to the ink injection. The velocity of ink transportation in the gut was calculated using kymograph images generated by ImageJ.

### Quantification of cloaca contractions

The region-of-interest (ROI) was set around a cloaca, and changes in movement of ROI in captured images were converted into intensity values using the Stack Difference of ImageJ. The values of intensity were normalized by the first frame of filmed data.

### Immunohistochemistry

Guts were fixed in 4% (w/v) paraformaldehyde (PFA)/phosphate buffered saline (PBS: 0.14 M NaCl, 2.7 mM KCl, 10 mM Na_2_HPO_4_–12H_2_O, 1.8 mM KH_2_PO_4_) overnight at 4 °C, and were washed in 0.1% Triton X-100 in PBS (PBST) three times for 5 min each at room temperature (RT). After blocking with 0.5 % blocking reagent (Roche, 1096176)/PBS overnight at 4 °C, the specimen was incubated overnight at 4°C with a 1:300 dilution of anti-Tuj1 (R & D systems, MAB1195) in 0.5 % blocking reagent/PBS. Following three times washing in PBST (each 1 h) at RT, the specimen was reacted with a 1:300 dilution of Alexa 568 (Invitrogen, A10042) goat anti-mouse IgG in 0.5 % blocking reagent/PBST overnight at 4 °C. After washing three times in PBST (each 1 hr) at RT, fluorescent images were obtained using the Leica MZ10 F microscope with the DS-Ri1 camera or Nikon A1R confocal microscope.

### Statistical analysis

The box plots represent the median, upper and lower interquartile. Student’s or Welch’s t-test were conducted using Excel to compare data statistically. Graphs were made by Excel or R.

## Results

### Bidirectional progression of peristaltic waves is dominant in embryonic guts (E8 to E12)

In this study, we have assessed the gut peristalsis by combining video recordings and kymography as described in the Materials and Methods. In brief, the entire gastrointestinal tract posterior to the duodenum is dissected from an embryo, and placed into a petri dish with pins to secure the specimen, whose hairpin curved structure at the umbilicus is retained with minimum artificial stress (Fig. 1A, B and Movie 1). Since the peristalsis is very sensitive to a mechano-stimulation such as a touch with forceps, the specimen is allowed to rest for 10 min, which is followed by the video recording for another 10 min in a way similar to the previous report (Chevalier, 2018). The filmed data are processed digitally using ImageJ (Fig. 1C), and subjected to kymography (Fig. 1D). If a gut undergoes regular peristalsis, for example one contraction per minute, the kymograph shows 10 repeated lines. Largely, two characteristic patterns are recognized: one is a line of inverted v-shape associated with an apex, and the other is simply a slanted line with no recognizable apex (Fig. 1E, F). The inverted v-shaped line shows bidirectionally propagating peristaltic waves with the apex corresponding to the origin of waves. The simply slanted lines likely represent a unidirectional wave propagation. As explained in more detail below, we have found that the majority of lines in the kymograph prepared from the midgut (developing intestines including jejunum and ileum) are of the v-shaped type accompanied by an apex (Fig. 1G; purple in the graph), suggesting that bidirectional peristalsis is dominant in embryonic intestines, at least from E8 to E12. Therefore, in further analyses we have been focusing on the origins of bidirectional peristaltic waves.

**Figure 1.**
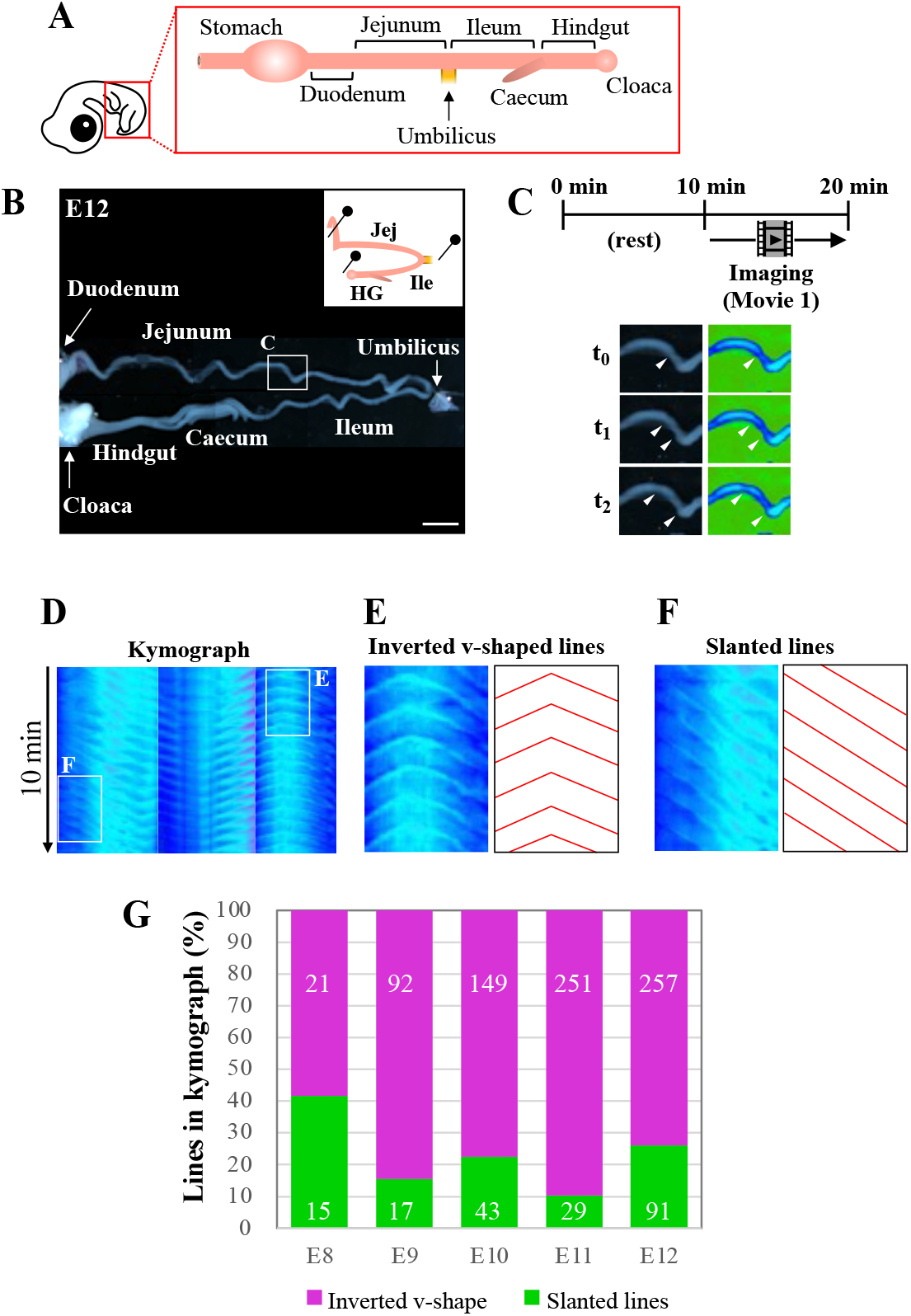
Kymography reveals that bidirectional waves are predominant in embryonic gut peristalsis, at least until E12. (A) The segment of gut located posteriorly to the stomach is dissected from an embryo, and (B) placed into a petri dish with gentle holding with pins to avoid dislocation of the specimen. (C) The specimen is video recorded for 10 min following 10 min of rest in a dish, and subsequently subjected to digital processing with ImageJ so that peristaltic movements can be visualized by changes in colors. The photos, showing an origin and the propagating waves, are three frames extracted from Movie 1 at the portion indicated in a square in B. (D) Digital images are further processed to make kymographs. Periodic propagation of peristaltic waves is represented by repeated lines along the time period of 10 min. (E, F) If the propagation of a wave is bidirectional which is accompanied by an origin, it is shown in the kymograph as an inverted v-shape with an apex (E), whereas unidirectionally propagating waves are represented simply by slanted lines without an apex (F). The majority of peristaltic waves are bidirectional in embryonic guts examined from E8 to E12. Scale bar: 4 mm.

We have measured the velocity of propagating waves in the jejunum and ileum at the stages from E8 and E12 (Fig. S1). Our observations are largely consistent with those reported previously (Chevalier et al., 2017) (see also Discussion).

### Distribution map of the origins of peristaltic waves

To determine how the origins of peristaltic waves (OPWs) emerge during the gut development, we have made a distribution map of the OPWs in the jejunum and ileum from E8 to E12. For each of these stages, 4 embryos were subjected to the video recording, digital processing, and kymography. Fig. 2A only includes one embryo per stage, while the other three repeats can be found in Fig S2. OPWs are demonstrated in single dots (Fig. 2a), which reflect the positions of apices associated with inverted v-shaped lines in the kymograph (see also Fig. 1E).

**Figure 2.**
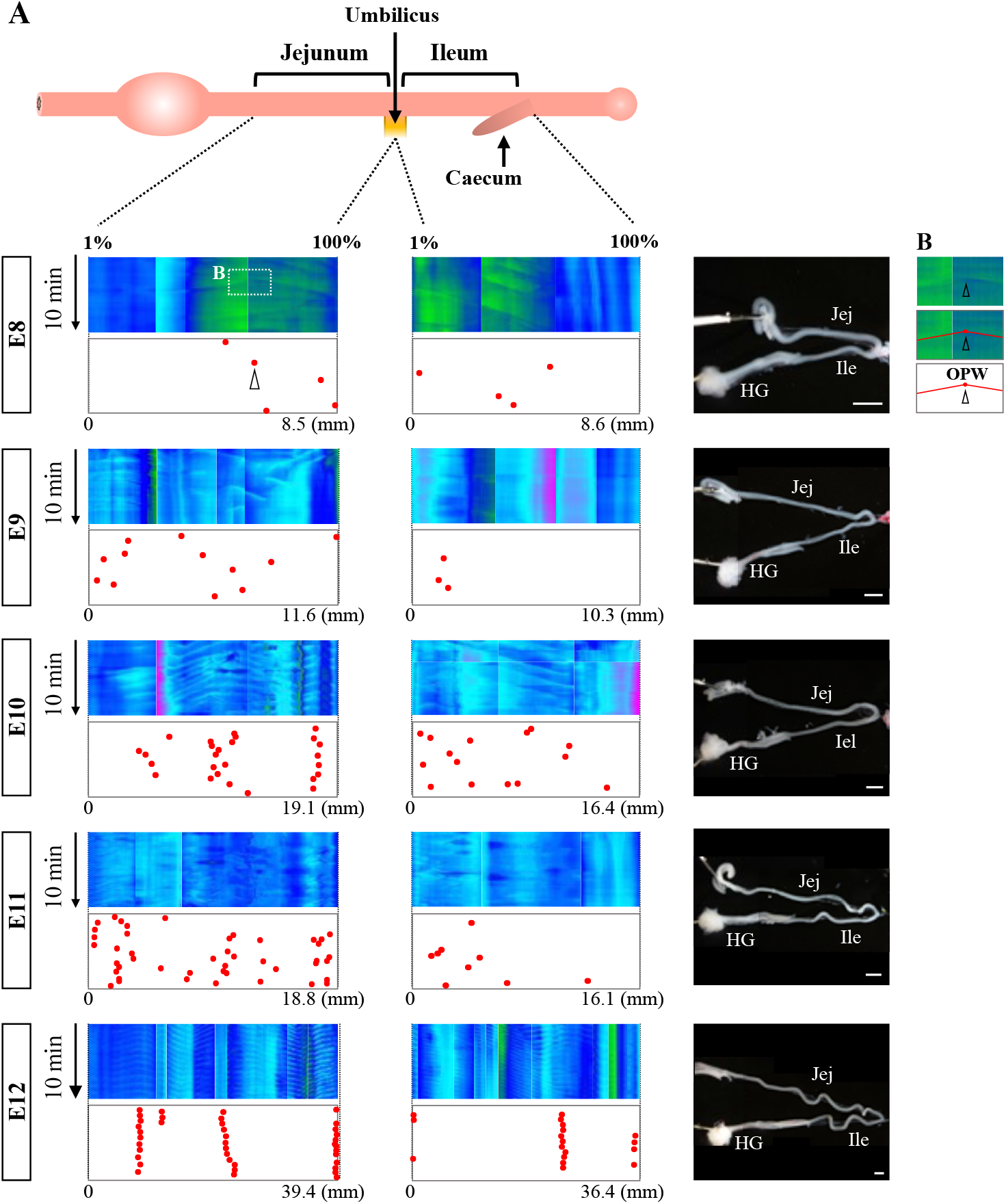
Distribution map of origins of bidirectional waves during gut development. (A) In the kymographs, the locations of apices are plotted to prepare the map of origins. For each stage, four embryos have been assessed and one of these is shown in this figure, whereas the other three are shown in Fig. S2. The locations of the origins are plotted in the jejunum and ileum. Since the length of the gut differs between individuals, the locations are normalized (1% to 100%) relative to the length of each region (see texts for details). While origins are randomly distributed at E8, they become confined to specific regions by E12. (B) The square indicated in the kymograph of E8 is enlarged to show the apex with v-shaped lines. Scale bars: 2 mm.

Since the gut is a long organ that keeps growing during development, kymography is performed separately in several different regions within the un-interrupted gut axis, and multiple kymographs are subsequently tiled (combined) so that the patterns of peristalsis in the entire gut can be readily assessed. Thus, abrupt changes in colors seen in a single kymograph panel along the gut axis are the artifacts of the tiling process. Another concern raised during these analyses is that the actual length of the gut varies among embryos even at the same developmental stage. To circumvent these problems, the jejunum and ileum are separately normalized in length, and location of OPWs is plotted as a relative position (1 % to 100 %). The umbilicus is a landmark for the border between jejunum and ileum, and the posterior end of the ileum is the site where the caeca protrude. For example, if an OPW is located at the border between jejunum and ileum, its position is either 100 % in the panel of jejunum, or 1 % of the ileum panel.

As revealed by the map (dots in Fig. 2A and Fig. S2), the positions of the OPWs become progressively stable from E8 onward. At E8, a few OPWs are randomly distributed, and by E11 the number of OPWs increases, although their distribution is still random. At E12, the OPWs are confined to several distinct positions: three major sites harbor the OPWs in the jejunum of embryo #1 (red dots in Fig. 2), meaning that once a given site starts to operate as an OPW, this site continues to serve as an origin of waves. 14.5 and 14.0 apices (average) at a single site are detected during 10 min in the jejunum and ileum (Fig. 2A, and Fig. 2S), meaning that waves occur at 24.2 mHz and 23.3 mHz frequency, respectively. Chevalier et al. reported 30 mHz frequency for the waves in the E12 jejunum (the frequency in the ileum was not described) (Chevalier et al., 2019). As will be discussed below, such differences might be caused by a different way of specimen preparations. Nevertheless, both studies agree that the frequency of peristalsis increases from E8 onward.

### The OPWs are confined to distinct zones along the midgut

We then asked whether the distribution pattern of the OPWs is determined stochastically or by some intrinsic rules such as genetic program. If the pattern is determined genetically, the distribution pattern would be expected to be conserved between individual embryos. To address this, we have overlaid the maps of the OPWs prepared from four different embryos for each of E8 to E12. Fig 3A demonstrates merged maps with #1 embryo (with red dots, shown in Fig. 2A), and #2 (blue), #3 (gray), #4 (green) shown in Fig. S2. We have noticed that dots in different colors progressively converge to distinct sites along the midgut, which is most evident at E12. For example, dots for four different embryos are aligned at the border between jejunum and ileum. Thus, the distribution pattern of the OPWs might obey some rules and not occur stochastically.

**Figure 3.**
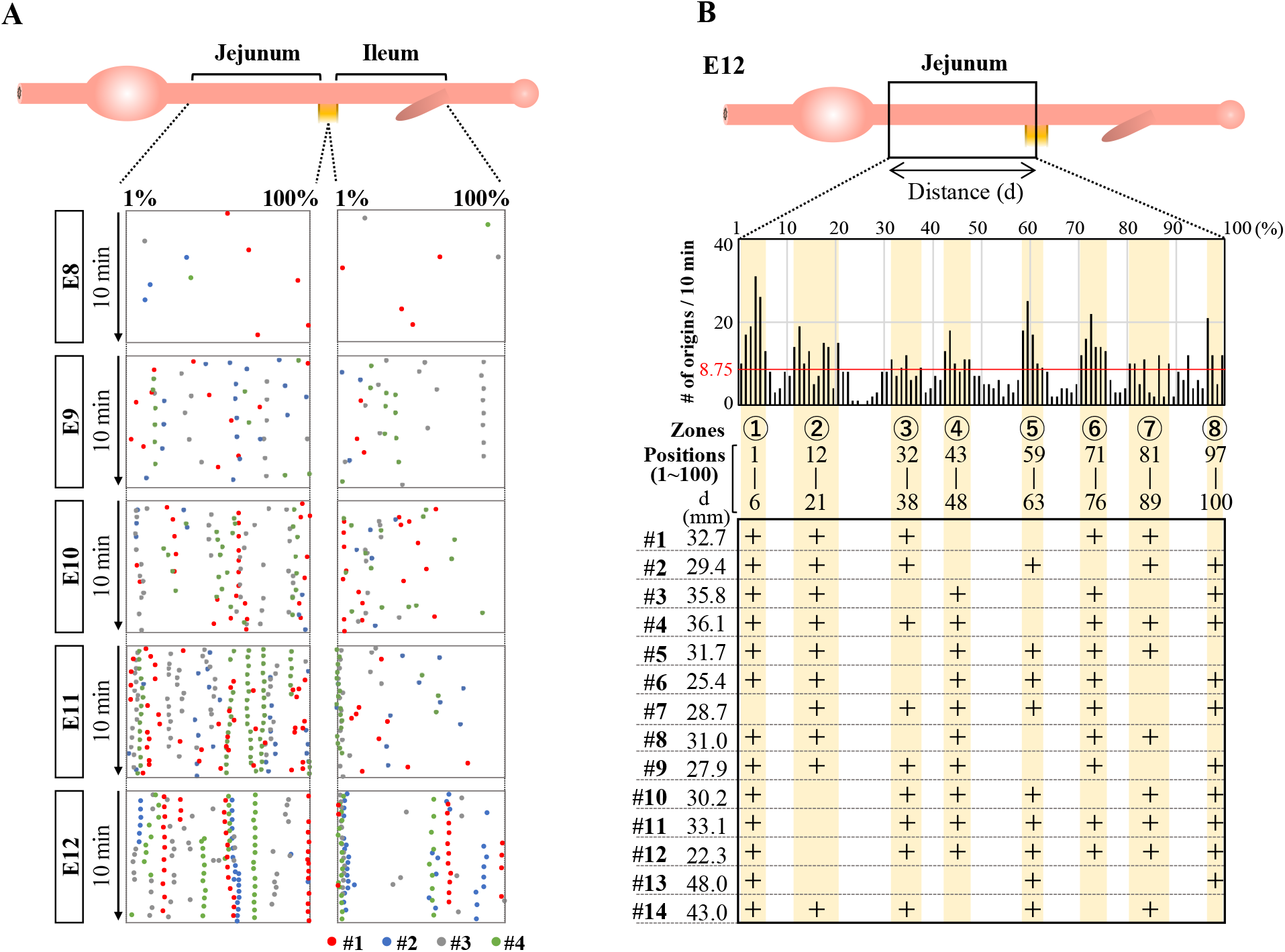
OPWs in the jejunum are progressively confined to distinct zones during gut development. (A) Distribution maps of four embryos are overlaid (#1 in red, Fig. 2; #2 to #4 as shown in Fig. S2). Dots in different colors progressively converge to distinct sites along the midgut. (B) Eight distinct zones (yellow) preferentially accommodate OPWs in the jejunum at E12. The total number of dots (OPWs) for 14 embryos is 875, and the average per one position is 8.75 (875/100; horizontal red line). The eight zones contain clusters of positions with each showing above 8.75. The zones allow two consecutive positions having below 8.75. The lower panel shows the usage of these zones in each embryo (#1 to #14). See text for details.

To further scrutinize this possibility, we have examined the distribution of the OPWs in the jejunum prepared from 14 different embryos at E12 (Fig. 3B, Fig. S3A). Since the actual length of the jejunum differs among embryos (22.3 mm to 48.0 mm), the position of the dots along the gut axis is normalized in the same way as shown in Fig. 2A and Fig. 3A, in which the length of the jejunum is divided into 100 discrete positions (1 to 100). The total number of the sites of OPWs (dots) for 10 min with the 14 specimens have been 875. If the distribution of the OPWs is perfectly stochastic, the 875 dots should be uniformly distributed with each position of 1 to 100 containing 8.75 dots (875/100 = 8.75; horizonal red line in the graph). The graph in Fig. 3B, however, shows that the distribution of OPWs is not uniform. We have noticed that dots are clustered in 8 different zones (shown in yellow) containing the dot number larger than 8.75 per position (above the red horizontal line), whereas other areas contain less than 8.75 per position. The position number in the 8 zones are as indicated in Fig. 3B. Thus, the zones 1-8 are the areas where OPWs are preferentially located. However, the embryonic jejunum of one individual does not necessarily use all the 8 zones. For example, zones 4, 5 and 8 are unused in #1 embryo, whereas # 13 embryo uses solely zones 1, 5, 8. Nevertheless, these findings suggest that zones 1-8 offer a favorable environment for the peristaltic waves to emerge.

### Enteric neural crest cells are required for the OPWs to be correctly patterned

It was previously shown that the ENS, which is derived from the neural crest, is not required for embryonic gut contraction (Chevalier, 2018; Chevalier et al., 2017) (Holmberg et al., 2007; Roberts et al., 2010). However, effects by ENS on the spatial distribution of the peristalsis were not examined. Therefore, we asked whether the correct distribution of the OPWs requires the ENS. Since there is no sign of correlation between the ENS pattern visualized by Tuj1 and the 8 zones (Fig S3B), we have decided to remove the ENS by ablating neural crest cells at the vagal level (somite1-7) of E1.5 embryo (Fig. 4), which is known to contribute to ENS in the jejunum (Fig. 4A) (Barlow et al., 2008; Burns and Douarin, 1998). Fig. 4B depicts the E12 jejuna of normal and neural crest-ablated embryos; Tuj1-staining of these jejuna shows little or no ENS in the neural crest-ablated gut.

**Figure 4.**
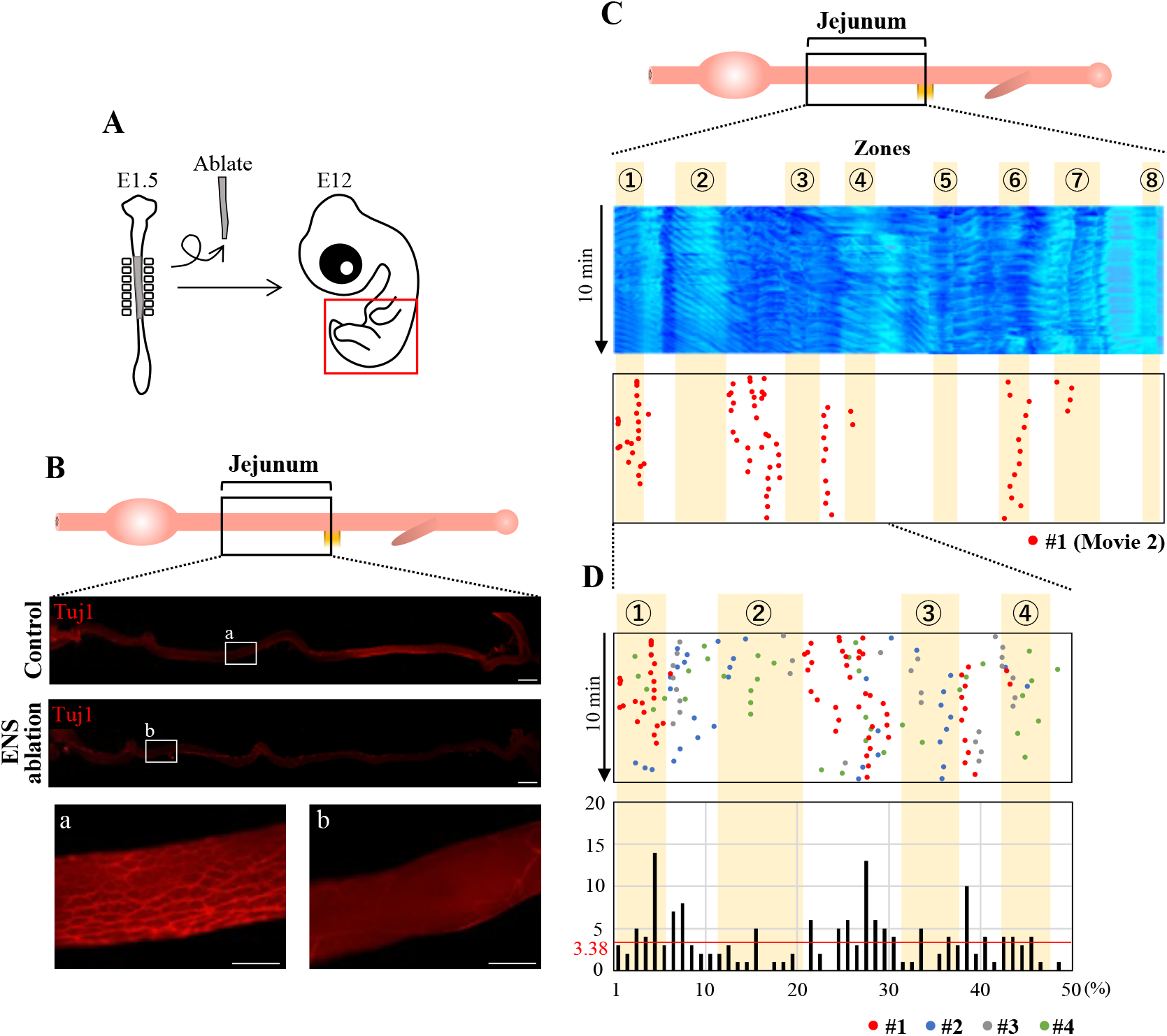
Enteric nervous system (ENS) is required for the correct distribution of OPW in the jejunum. (A) Neural crest cells that would normally contribute to the ENS are deprived by ablating the neural tube at the vagal level. (B) ENS-ablated jejunum at E12 stained with TuJ1: whereas meshwork structures of ENS are seen in control (non-ablated) specimen (a), ENS-ablated jejunum shows no or little ENS (b). (C) Kymograph and map of OPW distribution for the #1 specimen (red dots) of ENS-ablated jejunum. (D) Overlaid map for specimens #1 to #4 (red, blue, gray, green) in the anterior jejunum (50 positions), which would normally contain the four zones (1 to 4). The total number of OPWs is 169, and the average number is 3.38 (horizontal line in red). Many OPWs are found out-ofzone in an aberrant manner. Scale bars: 2 mm (low magnification) and 500 μm (high magnification).

Using the ENS-ablated jejunum dissected from four different embryos, we have assessed the positions of OPWs (Fig. 4C, Movie 2). To our surprise, several OPWs (dots) are found outside the previously identified zones. A merged map with four different embryos focused on zones 1 to 4 further highlights the aberrantly located OPWs (Fig. 4D). The total number of dots is 169, and the length of the jejunum is divided into 50 discrete positions. Thus, the average number of dots per position is 3.38 (169/50=3.38; horizontal red line). In the area between zones 2 and 3, many positions show an index larger than 3.38, with the largest being 13. Position 39, which is between zones 3 and 4 (Fig. 3B), also contains 10 dots. Thus, the ENS-ablation disrupts the distribution of the OPWs, suggesting that the enteric neural crest cells are important for the peristaltic waves to be correctly patterned along the gut axis.

### Neuronal activity is required for the correct distribution of the OPWs

How do the enteric neural crest cells regulate the positioning of the OPWs? Are cellular substances or neural activity (or both) important? To address these questions, we have immersed the entire jejunum (E12) in medium containing Na^+^ channel blocker tetrodotoxin (TTX), which blocks all neural activities. To accurately compare untreated and treaded specimens, an entire jejunum is prepared from an embryo and video-recorded for 10 min in normal medium. Following a 10 min rest, TTX is subsequently applied and a second recording is taken (Fig. 5A).

**Figure 5.**
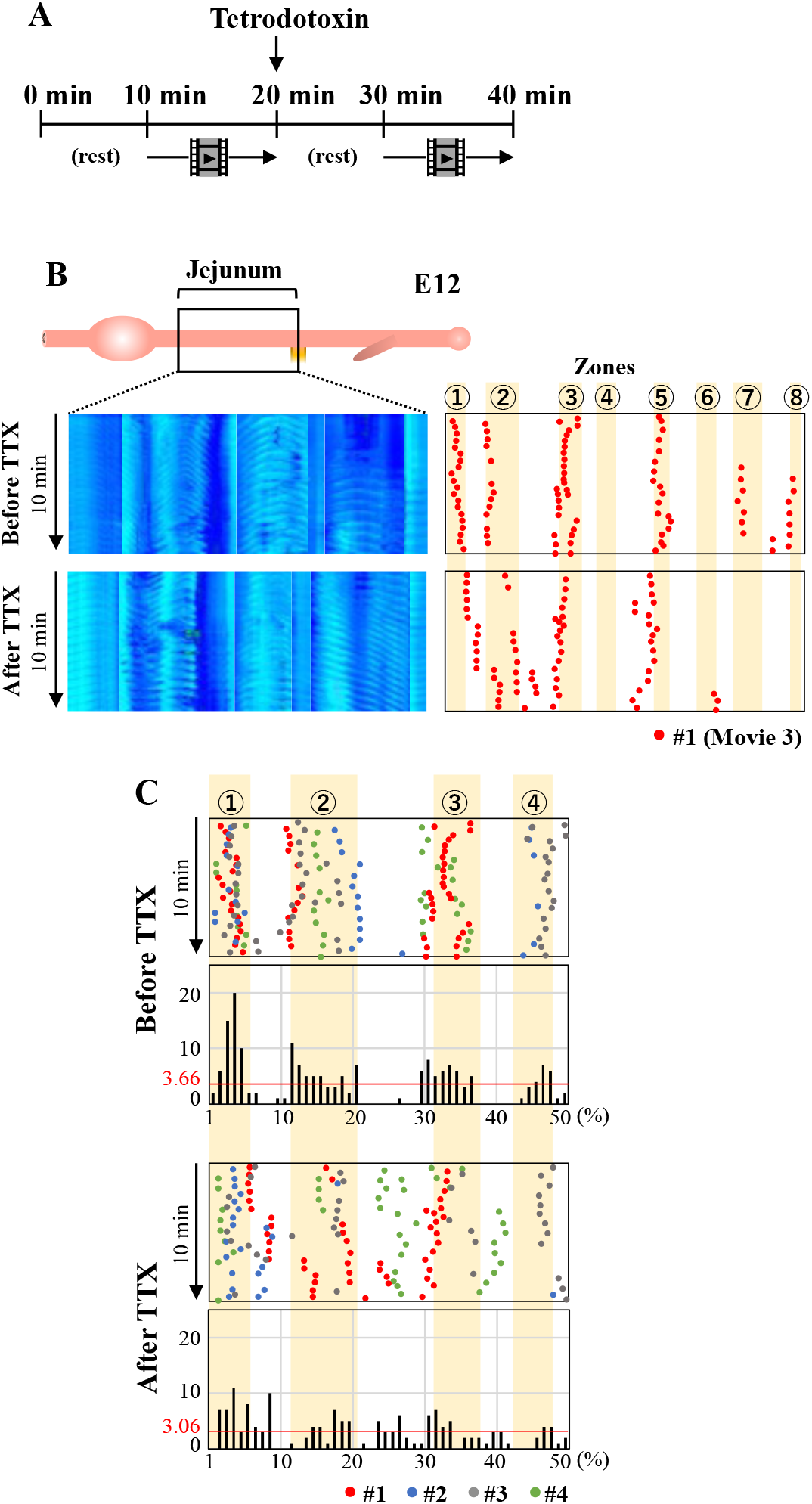
Blocking neural activity of ENS by TTX causes aberrant distribution of OPWs. Jejunum at E12 is immersed in a medium containing TTX. Jejunum is video-recorded in for 10 min in normal medium. (A) Following a 10 min rest, TTX is applied and a second recording is taken. (B) Kymographs and OPW distribution maps of a single jejunum (#1; red) before and after administration of TTX. (C) Overlaid map for specimens #1 to #4 (red, blue, gray, green) in the anterior jejunum (50 positions) in a manner similar to Fig. 4. The total number of OPWs is 187 and 153 for before and after TTX.

Fig. 5B and Movie 3 show the distribution of OPWs in embryo #1 (red dots). Before TTX administration, OPWs are found in 6 different zones. However, after the TTX treatment, 42 dots emerge outside the zones; 8 dots between zones1 and 2, 16 dots between zones 2 and 3, and 18 dots between zones 4 and 5. Fig. 5C shows a merged map with 4 different embryos for the region containing zones 1 to 4. Among the total of 153 dots identified during 10 min, 58 dots are located outside the zones, and 5 positions outside of these zones show more dots than the average of 3.06 (153/50 = 3.06). The total number of dots are comparable between untreated and TTX-treated specimens.

These findings suggest that neural activity by ENS is important for the correct distribution of the OPWs in the embryonic jejunum. Cellular factors elicited by differentiating neural crest cells might play a minor role, if any.

### ENS is required for the proper transportation of the inter-luminal contents

Does the perturbed OPW distribution have any impacts on the gut function? To address this question, we first examined whether the normal embryonic gut (E12) is capable of transporting inter-luminal contents by injecting fluorescent ink into the duodenum of the isolated gut (Fig. 6A). The fluorescent ink is successfully transported aborally. The velocity of the transportation varies between individuals; 1 cm/min to 39 cm/min (Fig. 6B, Movie 4). This is slightly unexpected because the majority of peristaltic waves are bidirectional at E12, as mentioned in Fig. 1. How the bidirectional waves enable the aboral translocation of inter-luminal contents remains unknown. We then conducted a negative control, in which fluorescent ink was injected into the gut immersed in Ca^2+^-free medium, conditions known to cease contraction (Chevalier et al., 2017). In these specimens, inter-luminal ink fails to move (0 cm/min, Fig. 6B, Movie 4).

**Figure 6.**
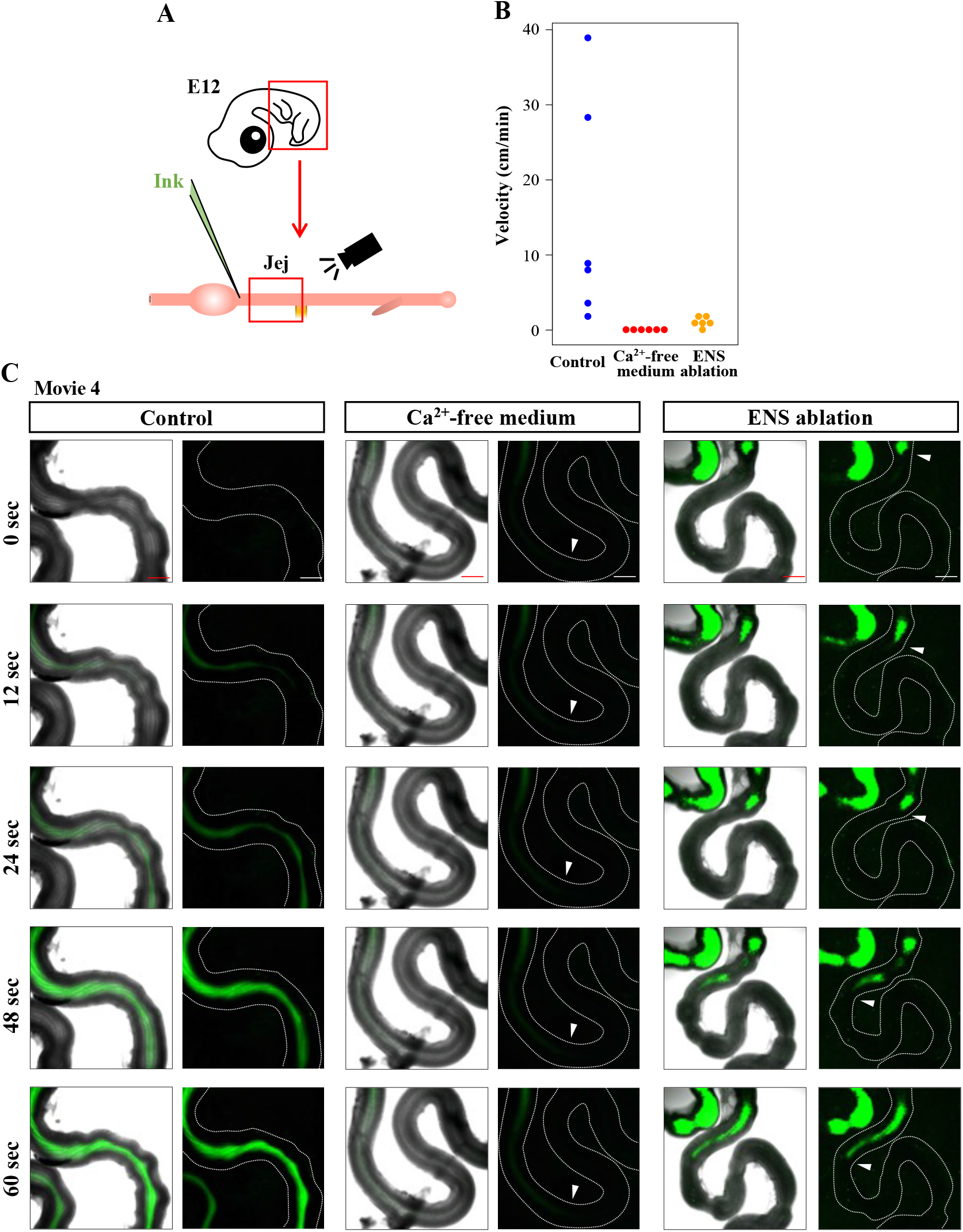
Requirement of ENS for smooth transportation of inter-luminal content in E12 jejunum. (A) Fluorescent ink is injected into duodenum using capillary, followed by video recording of jejunum. (B) Velocity of the injected ink in the jejunum. Each dot represents single jejunum during 6 repeats (embryos). Jejunums immersed in control and in Ca^2+^-free media, and ENS-ablated jejunum are assessed and compared. (C) Five frames are selected from each of movies (Movie 4). Arrowheads indicate a front position of the ink moving in an oral-to-aboral direction, although the ink transportation in the control jejunum is too fast to stay in a picture frame. Scale bars: 500 μm.

Finally, we examined ink transportation in the ENS-deprived gut (neural crest ablated), in which peristaltic contractions occur despite altered distribution of OPWs (Fig 4). Here, the inter-luminal ink showed very little transit (0 to 1 cm/min) (n=6, Fig. 6B, Movie 4). Collectively, these findings highlight that the correct pattern of the OPWs is crucial for the inter-luminal contents to be smoothly transported. The correct positioning of the OPWs might coordinate multiple peristaltic movements along the gut axis.

### Coordination between hindgut peristalsis and cloaca contractions

We have extended the analyses of peristalsis to the hindgut. Kymographs produced from E12 hindguts that contains the cloaca demonstrate repeated slanted lines which represent unidirectional waves propagating aborally with few, if any, bidirectional waves (first recording in Fig. 7A and Fig. 7B, Fig. S4 A). The cloaca is a posterior-end tissue which controls the excretion of the urine-feces complex as well as egg-laying (King and McLelland, 1981; Romanoff, 1960). Notably, we have observed acute contractions in the cloaca which occur repeatedly (Movie 5), and whose relative intensity could be quantified using ImageJ (Fig. 7B). Among 5 embryos examined, the average frequency of cloaca contractions is 4.6 during 10 min (Fig. S4C). Since these acute and strong contractions inevitably draw its connected hindgut, they are represented in the kymographs as horizontal lines in the hindgut (black lines in traced panels), which indeed perfectly match the pattern of spikes of cloaca contractions in the panel of relative intensity (Fig. 7B). Importantly, the horizontal lines in the kymograph are coupled with the end of slanted lines (95.7 %). These findings raise the interesting possibility that the acute contractions in the cloaca are triggered by the arrival of peristaltic waves in the posterior end of the hindgut.

**Figure 7.**
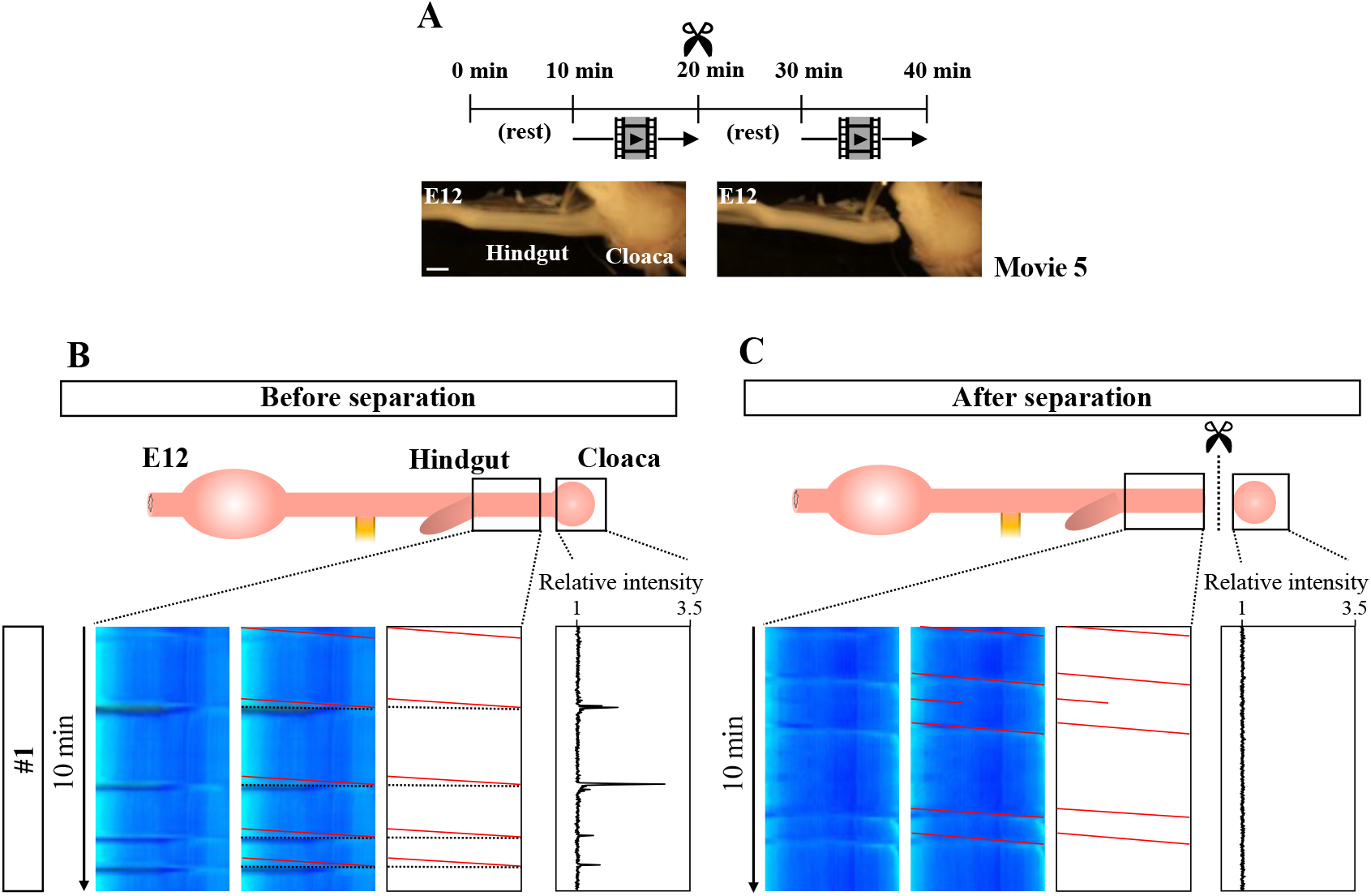
Tight correlation between peristalsis in the hindgut and cloaca’s strong contraction. (A) Experimental design. After the first video-recording of the cloaca-connected hindgut for 10 min, the cloaca (and its surrounding tissue) is separated from the hindgut. Following a 10 min rest, a second recording is taken. (B) Slanted lines downward to the right (red lines) represent peristaltic movements that recur unidirectionally (oral-to-aboral). These slanted red lines are frequently accompanied by horizontal lines (dotted). Since these horizontal lines are perfectly coincided with strong contraction of cloaca shown as relative intensity, these lines are consequences of stretching of hindgut being pulled by the cloaca. (C) Cloaca-disconnected hindgut continues to display the unidirectional peristaltic movements (red slanted lines). However, Horizontal lines have disappeared in the kymograph. Likewise, an isolated cloaca no longer implements strong contractions. The experiments have been repeated 5 times, and the other 4 are shown in Fig. S4. Scale bar: 1 mm.

To test this possibility, we have separated the cloaca (and its surrounding tissues) from the hindgut, and assessed the movements of these tissues. Similar to the procedure outlined in Fig. 5, the cloaca was separated after making a recording of the cloaca-connected hindgut (Fig. 7A) to precisely compare the contractions before and after the separation. Whereas the horizontal lines in kymographs have disappeared after the separation as expected, the peristaltic patterns in the hindgut are retained (Fig. 7B and C, Fig. S4B). Thus, the horizontal lines in the kymograph seen in the control specimen (cloaca-connected hindgut) are indeed attributed to the cloaca contractions. To our surprise, however, the isolated cloaca has completely ceased its acute contractions (Movie 5, Fig. 7C, Fig. S4B), indicating that the cloaca contractions take place in a hindgut-dependent manner. It is likely that the coordination between the hindgut and cloaca are mediated by the hindgut peristalsis, i.e. when a peristaltic wave reaches the posterior end of the hindgut, this triggers the acute contraction of cloaca.

In early embryos such as E8, both the hindgut peristalsis and cloacal contractions are seen, the latter being much weaker than those at E12. However, these two events occur independently (Fig. S4 D and Movie 6), suggesting that the coordination between the hindgut and cloaca is established later than E8.

### Bidirectional-to-unidirectional switch in the caecal waves from E8 to E12

Finally, we have examined the peristaltic patterns in the caeca. Unlike mammalians, the caeca in avians consist of a pair of long tubular structures protruding from the border between the midgut and hindgut, with the right and left portions of the caeca being morphologically symmetric (Fig. 8A). Conspicuous movements of peristalsis are detected along the caecal axis at E12 (Movie 7), consistent with previous reports (Chevalier et al., 2017). We have also observed overt peristalsis as early as at E8, which was not previously described. Kymographs prepared from the entire caeca at E8 or E12 demonstrate complicated sets of lines mixed with v-shaped (bidirectional), slanted (unidirectional), and horizontal lines (Fig. 8B, C). As explained in Fig. 7, the horizontal lines represent the drawing of the hindgut caused by the cloacal contractions. In both right and left caeca, whereas bidirectional waves are predominant at E8 (purple in Fig. 8B, D), unidirectional waves are more frequently seen at E12 caeca (green in Fig. 8C, D). The majority of these unidirectional waves propagate in a proximal-to-distal (P-to-D) direction (Fig. 8E, n=6 for each).

**Figure 8.**
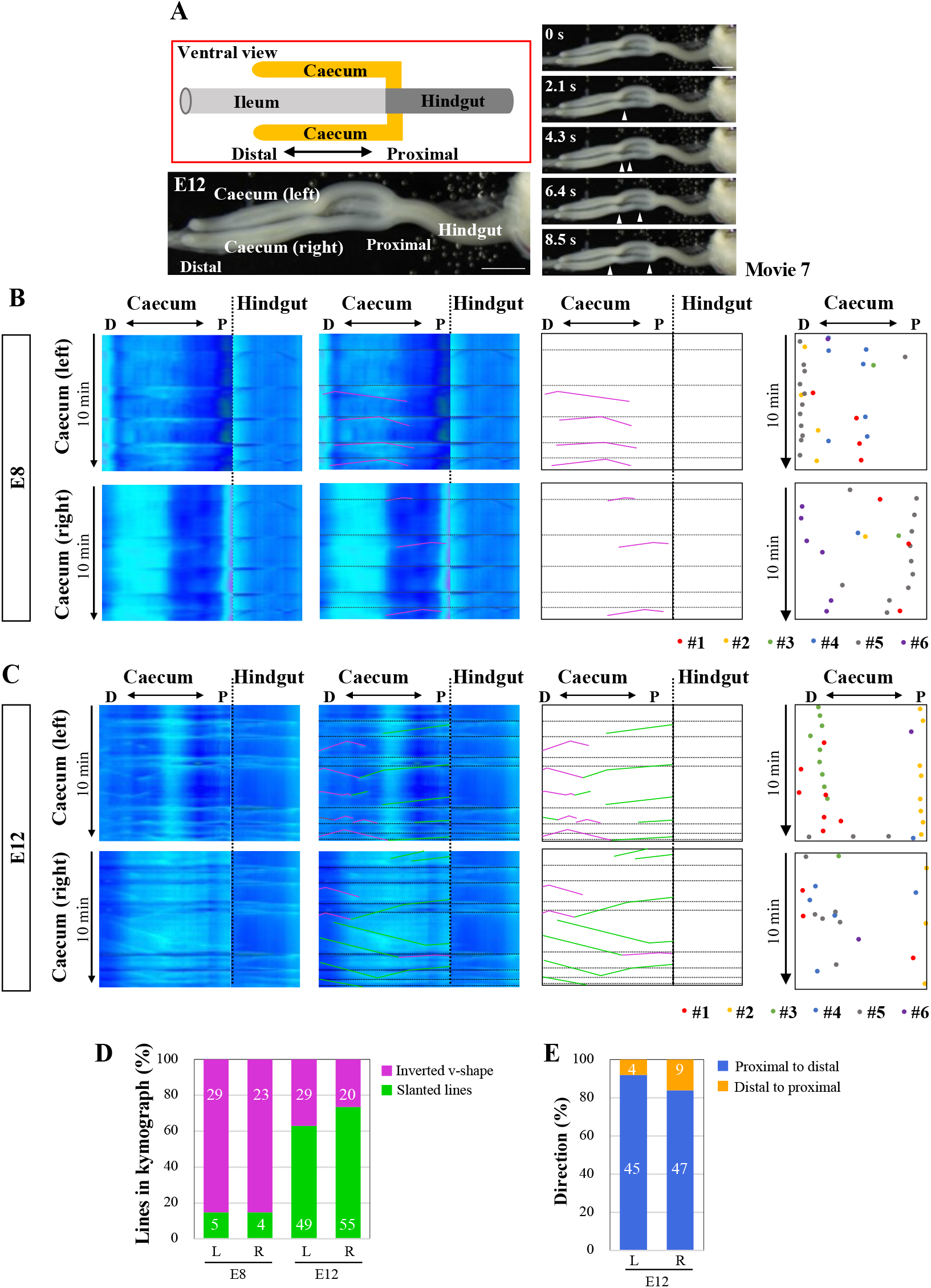
Peristaltic movements and OPW distribution map in the embryonic caeca. (A) Characteristic caecal structures in chicken embryos. Five frames are selected from Movie 7. (B) Peristaltic movements (v-shaped lines) are detected by E8 in both right and left arms. Horizontal lines are the continuation of those in the hindgut as explained in Fig. 7. No left-right (LR) symmetric patterns are found. (C) By E12, v-shaped (red) and simply slanted lies (green) are seen in kymographs. Horizontal lines are as explained in (C). Contrasting with the jejunum (Fig. 3), no zones of OPW confinement are detected in the caeca at E8 and E12 as shown in the distribution maps for 6 different embryos for each stage (B, C). (D) At E8, the majority of peristaltic movements are bi-directional (v-shaped lines with apex), whereas uni-directional movements (slanted lines) dominate later at E12. The number of lines found in the kymograph is shown in bars. (E) The majority of unidirectional movements at E12 are in a proximal-to-distal direction in both right and left arms. The number of lines found in the kymograph is shown in bars. Scale bars: 2 mm.

Similar to Fig. 2, a distribution map of the origins of bidirectional/peristaltic waves (OPWs) was prepared for each of right and left caeca at E8 and E12 (n=6 for each). Unlike those in the midgut, OPWs remain randomly distributed at E12, suggesting that OPWs emerge sporadically in the caeca at least until E12 (Fig. 8B, C). Despite the symmetrical morphology between the right and left (R-L) protrusions, the patterns of OPWs distribution and unidirectional wave propagations are not R-L symmetrical. Whether these caeca are able to transport inter-luminal contents is unknown.

To determine whether the peristaltic movements in the caeca are influenced by the main tract (midgut or hindgut), we have separated the caecal protrusions from the main tract at E12. To prevent an unnecessary displacement of the separated pieces, which would hamper the video recording and kymography, the mesentery that encapsules the gut is retained so that the separated caecum is secured within the mesentery (Fig. S5A, Movie 7). The patterns of OPWs are largely similar before and after the separation (Fig. S5B), suggesting that the peristaltic movements in the caeca take place autonomously at least until E12. A slight difference is observed for the velocity of wave propagation: P-to-D and D-to-P waves become slower and faster after the separation, respectively (Fig. S5C).

## Discussion

We have demonstrated the distribution patterns of peristaltic waves in the embryonic chicken gut including jejunum, ileum, hindgut, and caeca, by combining video recording and kymography. In the E8-E12 midgut (jejunum and ileum), the waves are predominantly bidirectional accompanied with an origin of waves which is detected as an apex on the kymographs. A distribution map of the origins of peristaltic waves (OPWs) on the kymographs reveals progressive stabilization of OPWs, and a requirement of neural activities of ENS for this stabilization during gut development. Furthermore, we have shown that peristaltic movements in the hindgut are coupled with cloaca contractions, highlighting inter-regional coordination along the gut axis.

A series of pioneering work by Chevalier and coworkers have described embryonic peristalsis mostly in chickens with a particular focus on local velocity, frequency and amplitude of peristaltic waves in given regions (Chevalier et al., 2020; Chevalier et al., 2019; Chevalier et al., 2017). The velocities we have observed in the current study (Fig. S1) are largely consistent with their reports, although some differences are noticed for the velocity at E9: 1.3 mm/min and 1.6 mm/min in the jejunum and ileum in our assessment, respectively, whereas 0.7 mm/min and 0.7 mm/min in (Chevalier et al., 2017). These differences might be attributed to how the gut specimens are placed in a petri dish. Chevalier et al. used a minimum amount of liquid medium so that the surface tension of medium could hold the gut, whereas in our assay the gut is immersed in an excessive amount of liquid medium to avoid the surface tension which would cause mechano-stimulation on the gut motility. In addition, in their study, a fragment of 1 cm long of jejunum at E12 was used to measure the velocity, whereas we have used a non-interrupted gut as explained above (Fig. 1B). Thus, our analyses have provided novel spatial information about the distribution of waves along the entire gut axis.

### Spatial stabilization of OPWs during gut development

In the developing midgut, OPWs which are randomly distributed at earlier stages become progressively stabilized by E12 until periodic and regular patterns form. The stabilized sites of OPWs are clustered into 8 distinct zones in the jejunum. These 8 zones are largely conserved between individuals, raising a possibility that the positions of the OPWs along the gut axis could be determined by genetic program, the details of which have yet to be identified. It is noteworthy that OPWs are frequently localized at the border between the jejunum and ileum, the site connected to the umbilicus. This raises a possibility that the zones of OPWs might be influenced by mechanical forces generated by, for example, a tubular branching point.

### Neural activity is required for the stabilization of OPWs and inter-luminal transportation in embryonic gut

Chevalier and coworkers reported previously that TTX administration did not affect local velocity, frequency or amplitude of peristaltic waves in given local regions or isolated pieces of embryonic chicken gut (Chevalier, 2018; Chevalier et al., 2019; Chevalier et al., 2017). The OPW distribution map produced in our study shows that neural activities are indeed not required for determining the frequency (number) of OPW emergence, since the number of OPWs is comparable between untreated-, ENS ablated-, and TTX immersed guts (Fig. 4, 5), consistent with previous reports (Chevalier et al., 2017; Holmberg et al., 2007; Roberts et al., 2010). However, our study has revealed that the neural activities play a pivotal role in the restriction OPWs to specific regions (zones 1 to 8) (Fig. 4, 5). Furthermore, the neural activity-regulated patterning of OPWs is functionally important: an ENS-deprived gut fails to smoothly translocate the inter-luminally injected fluorescent ink (Fig. 6). Whether this ineffective transportation is attributed solely to the disrupted spatial patterning of OPWs, or whether other factors stemmed by ENS-ablation are involved is currently unknown.

### Inter-region coupling of peristaltic contractions along the gut axis

We have demonstrated that the arrivals of the peristaltic waves to the posterior end of the hindgut are coupled with acute contractions of cloaca, suggesting intimate coordination between these tissues. In addition, an isolated cloaca ceases its own contractions, highlighting a requirement of hindgut-derived signals. Although precise mechanisms are currently unknown, it is conceivable that the hindgut muscles at the posterior end that are electrically activated upon the arrival of aborally propagating peristaltic wave might transmit signals to the coprodaeum (the innermost division of the cloaca) through intercellular signaling, e.g., gap junctions. Such gap junctions couple the smooth muscles that effect the peristaltic movements (Chevalier et al., 2017).

Contrary to the cloaca, the caecum appears to undergo peristalsis in an autonomous manner, since the peristalsis does not change upon a separation from the main gut tract. The avian caecum is unique and composed of a pair of long tubular structures. It is thought that this characteristic structure is important for the process of the urine and feces: unlike mammals, contents in the colon are back-transported to the caecum where they are further processed so that any residual nutrients and water are thoroughly absorbed (Clench, 1999; Clench and Mathias, 1995; Duke, 1989). Thus, the caecal specific peristalsis might be important for this efficient processing of internal contents. Unveiling the mechanisms by which the peristalsis is regulated in the caecum in avians is expected to help understanding the caecum in mammals including humans, the function of which remains largely unexplored.

Lastly, defects in peristaltic movements are associated with a variety of disorders in the gastrointestinal tract (Boeckxstaens et al., 2016; Camilleri et al., 2008; Sanders et al., 2006; Schneider et al., 2019), and development of therapeutic treatments to ameliorate the peristaltic defects has long been awaited. Toward this goal, unveiling the cellular basis of gut peristalsis is critical. The chicken embryonic gut serves as a simplified experimental system enabling high resolution analyses because they undergo peristalsis even before hatching (no food contents in the gut). Chevalier et al. recently reported that the activity of gut peristalsis becomes higher at E12 onward, where networks between interstitial cells of Cajal, ENS and smooth muscles may become mature (Chevalier et al., 2020). Although analyzing guts post E12 is currently beyond our capabilities due to gut’s massive increase in length, it is important to know how the spatial patterns of peristalsis evolve and mature during late development.

## Supporting information

Moview 1 to 7

## Acknowledgements

We thank Hana Godfrey and Dr. Scott Gilbert for careful reading of the manuscript and discussion. This work was supported by AMED (JP17gm0610015), and JSPS KAKENHI Grant Numbers; 20K21425, 20H03259, 19H04775. Y.S is a fellow of JSPS.

## Supplementary Figures

**Fig. S1.**
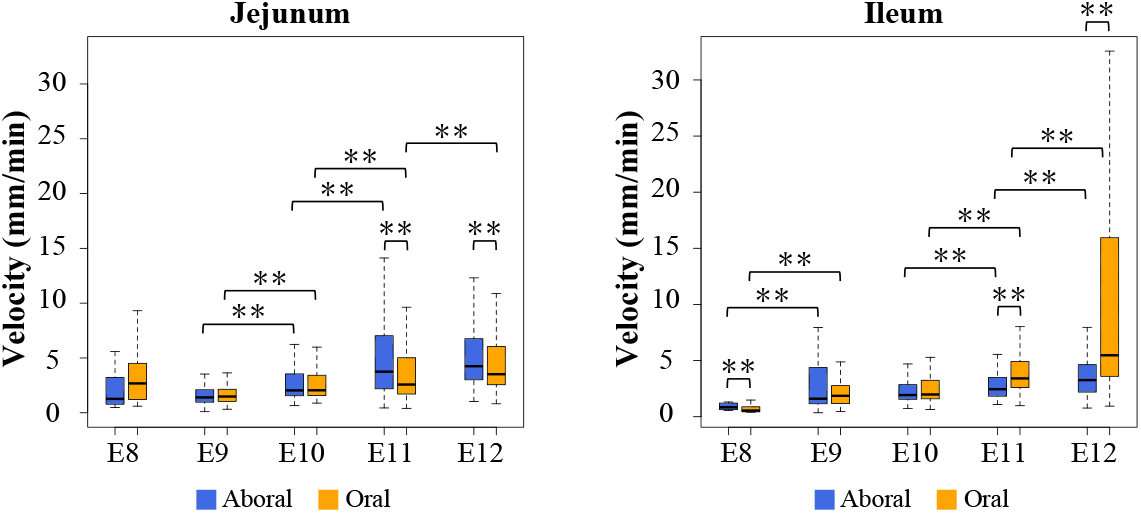
Velocity of propagation waves in the jejunum and ileum during gut development. Oral-to-aboral and aboral-to-oral movements are separately measured. **: p < 0.01. All the lines in the kymograph prepared from 4 embryos were subjected to calculation.

**Fig. S2.**
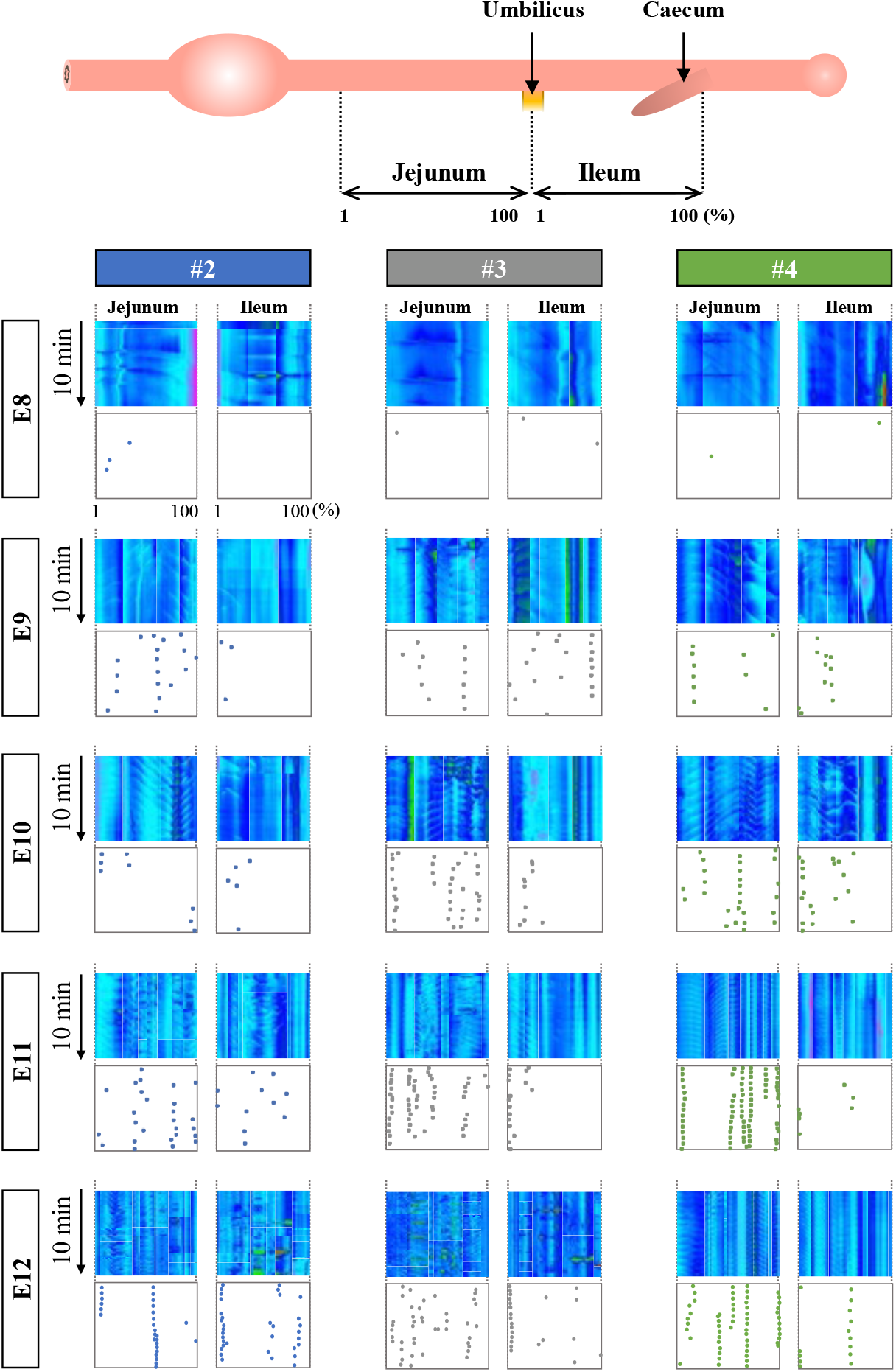
Same analyses explained in Fig 2. Individual embryos #2 to #4 are separately shown.

**Fig. S3.**
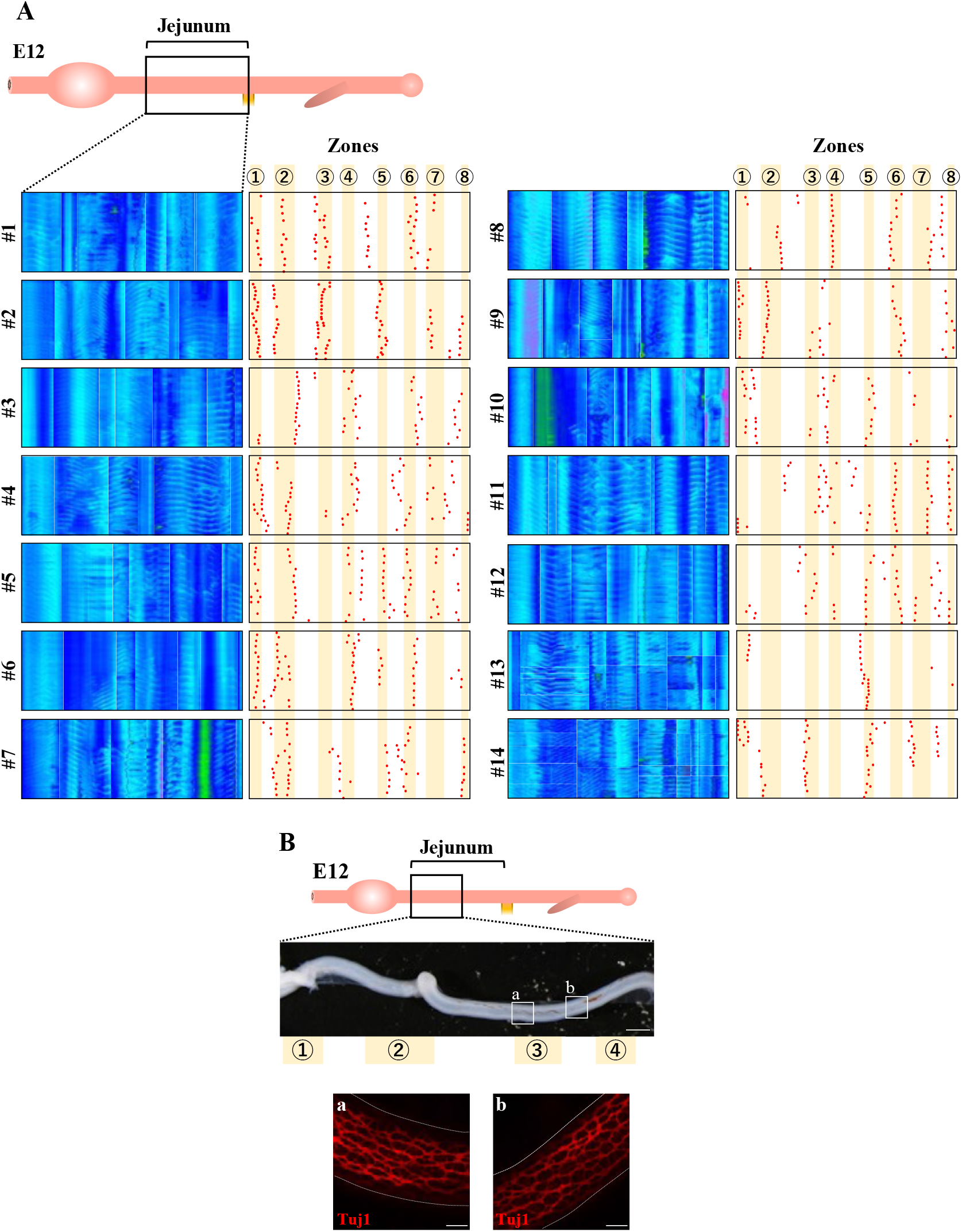
(A) Same analyses explained in Fig. 3B. Kymographs and OPW distribution maps are shown for each of 14 individuals. Zones 1 to 8 are as explained in Fig 3B and text. (B) Whole-mount staining with Tuj1 to visualize ENS in zone 3 and the inter-zone between zone 3 and 4. Scale bars: 2 mm (low magnification) and 200 μm (high magnification).

**Fig. S4.**
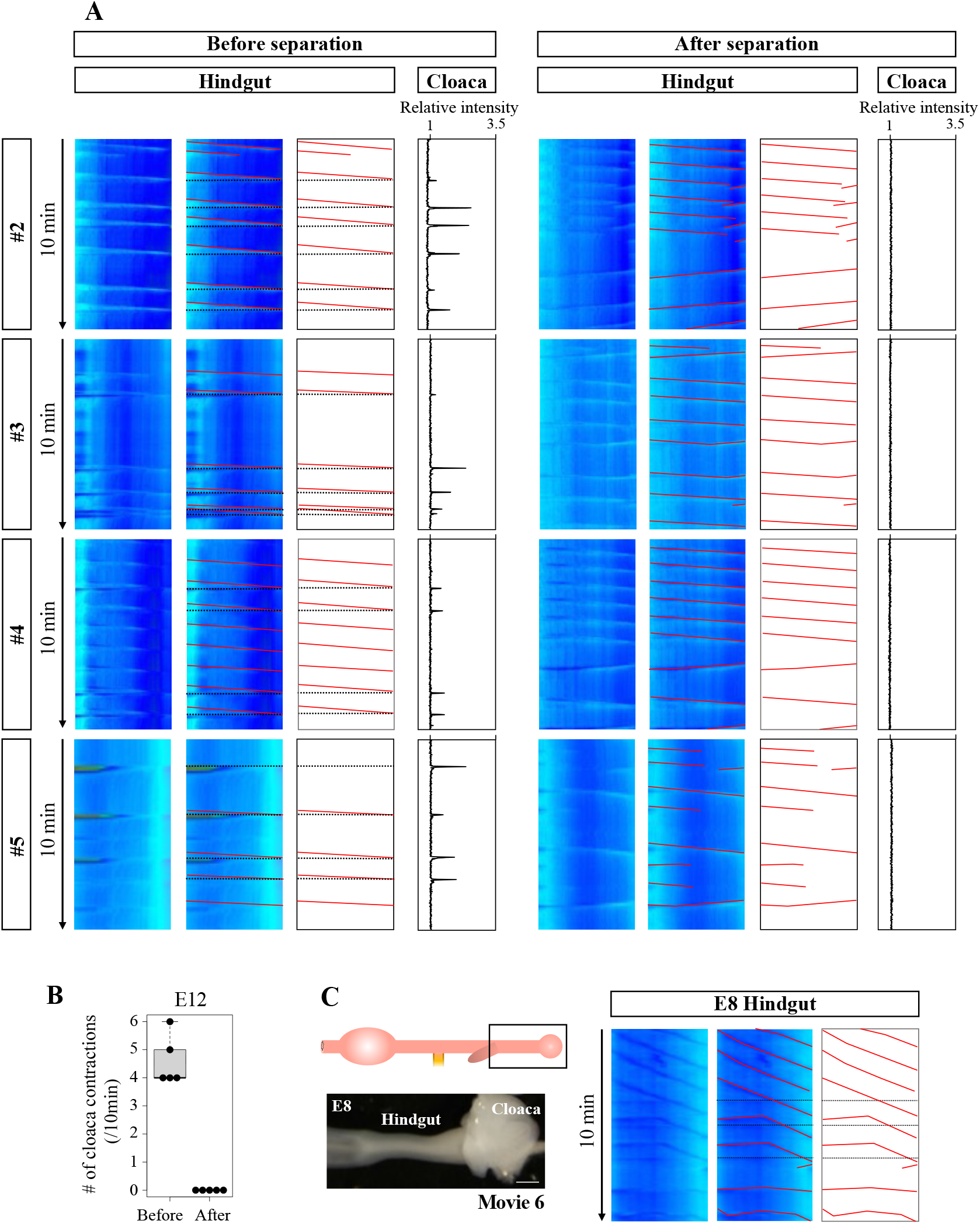
(A) Same analyses explained in Fig. 7. Four out of the five specimens are shown. (B) Number of cloaca contractions before and after the separation of cloaca. (C) Kymograph for the hindgut at E8. Peristalsis is not correlated with cloaca-derived horizontal lines. Scale bar: 500 μm.

**Fig. S5.**
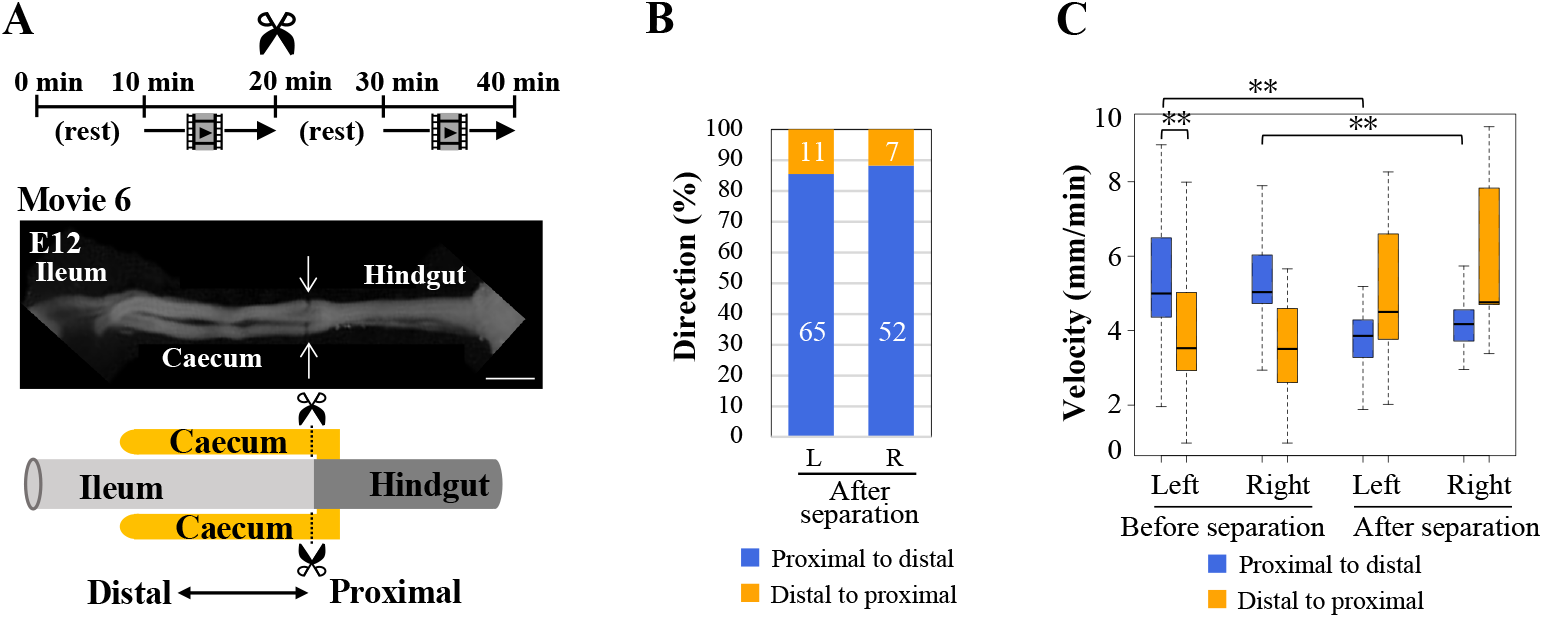
Isolation of the caeca from the main gut tract at E12. (A) The caeca are separated at their proximal ends, and video-recorded. (B) The proximal-to-distal predominance of peristalsis is retained in both arms. (C) Velocity of peristalsis is slightly affected by the separation in a direction dependent manner: slower and faster in a proximal-to-distal and distal-to-proximal directions, respectively. **: p < 0.01, scale bar: 2 mm.

**Movie 1: Bidirectional propagation of peristaltic waves in the jejunum at E12.**

Time-lapse images were obtained 2 frames per second. A white arrow shows an origins of peristaltic waves (OPWs) of bidirectional propagation. This video corresponds to Figure 1C.

**Movie 2: Aberrant distribution of OPWs in ENS-ablated jejunum at E12.**

Time-lapse images were obtained 2 frames per second. Red arrows show OPWs which emerge at ectopic sites between zone 2 and zone 3. This video corresponds to Figure 4C.

**Movie 3: Aberrant distribution of OPWs in TTX-treated jejunum at E12.**

Time-lapse images were obtained 2 frames per second before (control) and after TTX administration. White and red arrows are OPWs emerging inside and outside zones, respectively. This video corresponds to Figure 5B.

**Movie 4: Aberrant transportation of inter-luminally injected ink into ENS-ablated jejunum at E12.**

Time-lapse images were obtained 2 frames per second. Movements of inter-luminally injected fluorescent ink were recorded in jejunums immersed in control and in Ca^2+^-free media, and an ENS-ablated jejunum. This video corresponds to Figure 6C.

**Movie 5: Coupled contractions between hindgut and cloaca at E12.**

Time-lapse images were obtained 2 frames per second. White arrows show acute contractions of cloaca. The movie of cloaca-connected hindgut is followed by cloaca-disconnected hindgut of the same specimen. This video corresponds to Figure 7A.

**Movie 6: Un-coupled contractions between hindgut and cloaca at E8.**

Time-lapse images were obtained 2 frames per second. White arrow shows mild contractions of cloaca which are not coupled with peristaltic propagations in the hindgut. This video corresponds to Figure S4C.

**Movie 7: Peristaltic movements in the caeca at E8 and E12.**

Time-lapse images were obtained 2 frames per second. White arrows show directions of wave propagations. This video corresponds to Figure 8A.

## Notes

### Competing Interest Statement

The authors have declared no competing interest.

## References

Barlow, A.J., Wallace, A.S., Thapar, N., Burns, A.J., 2008. Critical numbers of neural crest cells are required in the pathways from the neural tube to the foregut to ensure complete enteric nervous system formation. Development 135, 1681–1691.

Bauer, A.J., Boeckxstaens, G.E., 2004. Mechanisms of postoperative ileus. Neurogastroenterol Motil 16 Suppl 2, 54–60.

Boeckxstaens, G., Camilleri, M., Sifrim, D., Houghton, L.A., Elsenbruch, S., Lindberg, G., Azpiroz, F., Parkman, H.P., 2016. Fundamentals of Neurogastroenterology: Physiology/Motility - Sensation. Gastroenterology.

Bornstein, J.C., Costa, M., Grider, J.R., 2004. Enteric motor and interneuronal circuits controlling motility. Neurogastroenterol Motil 16 Suppl 1, 34–38.

Bouchoucha, M., Devroede, G., Dorval, E., Faye, A., Arhan, P., Arsac, M., 2006. Different segmental transit times in patients with irritable bowel syndrome and “normal” colonic transit time: is there a correlation with symptoms? Tech Coloproctol 10, 287–296.

Burns, A.J., Douarin, N.M., 1998. The sacral neural crest contributes neurons and glia to the post-umbilical gut: spatiotemporal analysis of the development of the enteric nervous system. Development 125, 4335–4347.

Camilleri, M., McKinzie, S., Busciglio, I., Low, P.A., Sweetser, S., Burton, D., Baxter, K., Ryks, M., Zinsmeister, A.R., 2008. Prospective study of motor, sensory, psychologic, and autonomic functions in patients with irritable bowel syndrome. Clin Gastroenterol Hepatol 6, 772–781.

Chevalier, N.R., 2018. The first digestive movements in the embryo are mediated by mechanosensitive smooth muscle calcium waves. Philos Trans R Soc Lond B Biol Sci 373.

Chevalier, N.R., Ammouche, Y., Gomis, A., Teyssaire, C., Barbara, P.D., Faure, S., 2020. Shifting into high gear: how interstitial cells of Cajal change the motility pattern of the developing intestine. American Journal of Physiology-Gastrointestinal and Liver Physiology 319, G519–G528.

Chevalier, N.R., Dacher, N., Jacques, C., Langlois, L., Guedj, C., Faklaris, O., 2019. Embryogenesis of the peristaltic reflex. J Physiol 597, 2785–2801.

Chevalier, N.R., Fleury, V., Dufour, S., Proux-Gillardeaux, V., Asnacios, A., 2017. Emergence and development of gut motility in the chicken embryo. PLoS One 12, e0172511.

Clench, M.H., 1999. The avian cecum: Update and motility review. Journal of Experimental Zoology 283, 441–447.

Clench, M.H., Mathias, J.R., 1995. The Avian Cecum.

Duke, G.E., 1989. Relationship of cecal and colonic motility to diet, habitat, and cecal anatomy in several avian species. J Exp Zool Suppl 3, 38–47.

Hamburger, V., Hamilton, H.L., 1951. A series of normal stages in the development of the chick embryo. J Morphol 88, 49–92.

Holmberg, A., Olsson, C., Hennig, G.W., 2007. TTX-sensitive and TTX-insensitive control of spontaneous gut motility in the developing zebrafish (Danio rerio) larvae. J Exp Biol 210, 1084–1091.

Huizinga, J.D., Ambrous, K., Der-Silaphet, T., 1998. Co-operation between neural and myogenic mechanisms in the control of distension-induced peristalsis in the mouse small intestine. J Physiol 506 (Pt 3), 843–856.

Huizinga, J.D., Lammers, W.J., 2009. Gut peristalsis is governed by a multitude of cooperating mechanisms. Am J Physiol Gastrointest Liver Physiol 296, G1–8.

King, A.S., McLelland, J., 1981. Form and function in birds. Academic Press, London.

Mattei, P., Rombeau, J.L., 2006. Review of the pathophysiology and management of postoperative ileus. World J Surg 30, 1382–1391.

Pimentel, M., Soffer, E.E., Chow, E.J., Kong, Y., Lin, H.C., 2002. Lower frequency of MMC is found in IBS subjects with abnormal lactulose breath test, suggesting bacterial overgrowth. Dig Dis Sci 47, 2639–2643.

Roberts, R.R., Ellis, M., Gwynne, R.M., Bergner, A.J., Lewis, M.D., Beckett, E.A., Bornstein, J.C., Young, H.M., 2010. The first intestinal motility patterns in fetal mice are not mediated by neurons or interstitial cells of Cajal. J Physiol 588, 1153–1169.

Romanoff, A.L., 1960. The Avian Embryo. Macmillan, New York.

Sanders, K.M., Koh, S.D., Ward, S.M., 2006. Interstitial cells of cajal as pacemakers in the gastrointestinal tract. Annu Rev Physiol 68, 307–343.

Schneider, S., Wright, C.M., Heuckeroth, R.O., 2019. Unexpected Roles for the Second Brain: Enteric Nervous System as Master Regulator of Bowel Function. Annu Rev Physiol 81, 235–259.

Spencer, N.J., Dinning, P.G., Brookes, S.J., Costa, M., 2016. Insights into the mechanisms underlying colonic motor patterns. J Physiol 594, 4099–4116.

Takase, Y., Tadokoro, R., Takahashi, Y., 2013. Low cost labeling with highlighter ink efficiently visualizes developing blood vessels in avian and mouse embryos. Dev Growth Differ 55, 792–801.

